# Ing4-deficiency enhances HSC quiescence and confers resistance to inflammatory stress

**DOI:** 10.1101/2021.07.14.452366

**Authors:** Zanshé Thompson, Georgina A. Anderson, Melanie Rodriguez, Seth Gabriel, Vera Binder, Alison M. Taylor, Katie L. Kathrein

## Abstract

Hematopoiesis is tightly regulated by a network of transcription factors and complexes that are required for the development and maintenance of hematopoietic stem cells (HSCs). We recently identified the tumor suppressor, Ing4, as a critical regulator of HSC homeostasis. Though the Ing4 mechanism of action remains poorly characterized, it has been shown to promote stem-like cell characteristics in malignant cells. This activity is, in part, due to Ing4 mediated regulation of several major signaling pathways, including NF-κB and c-Myc. In murine hematopoiesis, Ing4 deficiency induces G_0_ arrest in HSCs, while simultaneously promoting gene expression signatures associated with differentiation. This results in a poised state for Ing4-deficient HSCs. Long term HSCs are unable to overcome this block, but short-term HSCs convert the poised state into regenerative capacity during hematopoietic challenges, including irradiation and transplantation. Overall, our findings suggest that Ing4 plays a crucial role in the regulation of hematopoiesis. Our model provides key tools for further identification and characterization of pathways that control quiescence and differentiation in HSCs.

## Introduction

Hematopoiesis is the process by which hematopoietic stem cells give rise to all mature blood cells ^1^. Hematopoietic stem cells (HSCs) simultaneously maintain quiescence and promote differentiation through tightly regulated cell-extrinsic and cell-intrinsic frameworks of proteins that both maintain, promote, and repress specific gene expression patterns to enable ^2–6^. This delicate balance achieves a constant source for immune cell production while retaining a robust stem cell pool ^7–14^. Disruption of the signaling pathways that maintain the stem cell pool can have significant consequences on the longevity and robust responses of the immune system. The importance of understanding hematopoiesis is appreciated clinically, as hematopoietic stem and progenitor cells (HSPCs) are used for bone marrow transplantation to treat a variety of diseases, including anemia, B-thalassemia, and inherited metabolic disorders ^15^.

To uncover critical regulators of HSCs, we recently identified Inhibitor of growth 4 (Ing4) as a regulator of HSC specification in a zebrafish reverse genetic screen ^12^. Ing4 is a tumor suppressor protein with frequently identified somatic mutations in human malignancies and Ing4 deficiency is associated with a poor prognosis in several different cancer types ^16–18^. Ing4 has many regulatory roles, both as a chromatin remodeling protein as a member of the Hbo1 complex and as a direct regulator of several major signaling pathways: NF-κB, c-Myc, p53, and HIF-1α ^19–25^.

Previous work on Ing4-deficiency showed that Ing4^−/−^ macrophages and neutrophils have increased Nκκ target gene expression and more cytokine production, suggesting a role for Ing4 in differentiated immune cell responses ^25^. Ing4 binds to the p65/RelA subunit of NF-κB, inhibiting p65 from binding to DNA and suppressing NF-κB target genes and inflammatory pathways ^21–24^. Ing4 has also been shown to directly bind and inhibit the c-Myc inhibitor, AUF1 ^26^, the HIF-1α inhibitor, HPH-2 ^27^, and enhance p53 through acetylation ^28^. These studies suggest that Ing4 is a critical regulatory protein, yet its role in hematopoiesis remains uncharacterized.

In this study, we identified an unexpected role for Ing4 as a negative regulator of HSC quiescence and self-renewal. Using an Ing4-deficient mouse model ^25^, we show that Ing4 deficiency disrupts normal HSC function. Ing4^−/−^ HSCs display increased quiescence with decreased sensitivity to chemotoxic insult and irradiation. Strikingly, loss of Ing4 expression has dueling outcomes for HSCs; they become more quiescence while simultaneously upregulating pathways associated with differentiation. While increased NF-κB mediated cytokine expression is observed in differentiated bone marrow cells, but not HSCs, this increase in cytokine expression does not impair ST-HSC function, unlike other models of NF-κB overexpression ^29,30^. c-Myc target genes including ribosomal biogenesis genes and genes associated with oxidative phosphorylation are also highly upregulated. Inhibition of c-Myc activity forces Ing4^−/−^ HSCs into cycling, but also results in reduced cell numbers. Our findings display an essential role for Ing4 in stem cell maintenance and differentiation.

## Results

### The Hematopoietic Program is Disrupted in the Absence of Ing4

To investigate the role of Ing4 in hematopoiesis, we used an Ing4-deficent mouse model to profile hematopoietic stem cells in the absence of Ing4 ^25^. When compared with wildtype (WT) mice, there was no significant difference observed in the percentage and absolute number of Lin^−^Sca1^+^c-Kit^+^ cells (LSKs) observed in the whole bone marrow (WBM) of Ing4^−/−^ mice (Figure 1a). Recent publications have identified two distinct populations of hematopoietic stem cells: quiescent, repopulating long-term HSCs (LT-HSCs), and more actively dividing short-term HSCs (ST-HSCs) ^4,31–34^. Further analysis of these specific HSC populations showed that, in the Ing4^−/−^ mice, the percentages of LT-HSCs (LSK CD34^−^CD150^+^) and ST-HSCs (LSK CD34^+^CD150^+^) were significantly increased (Figure 1d,e). These results suggest Ing4 deficiency may result in a differentiation block in HSCs.

**Figure 1.**
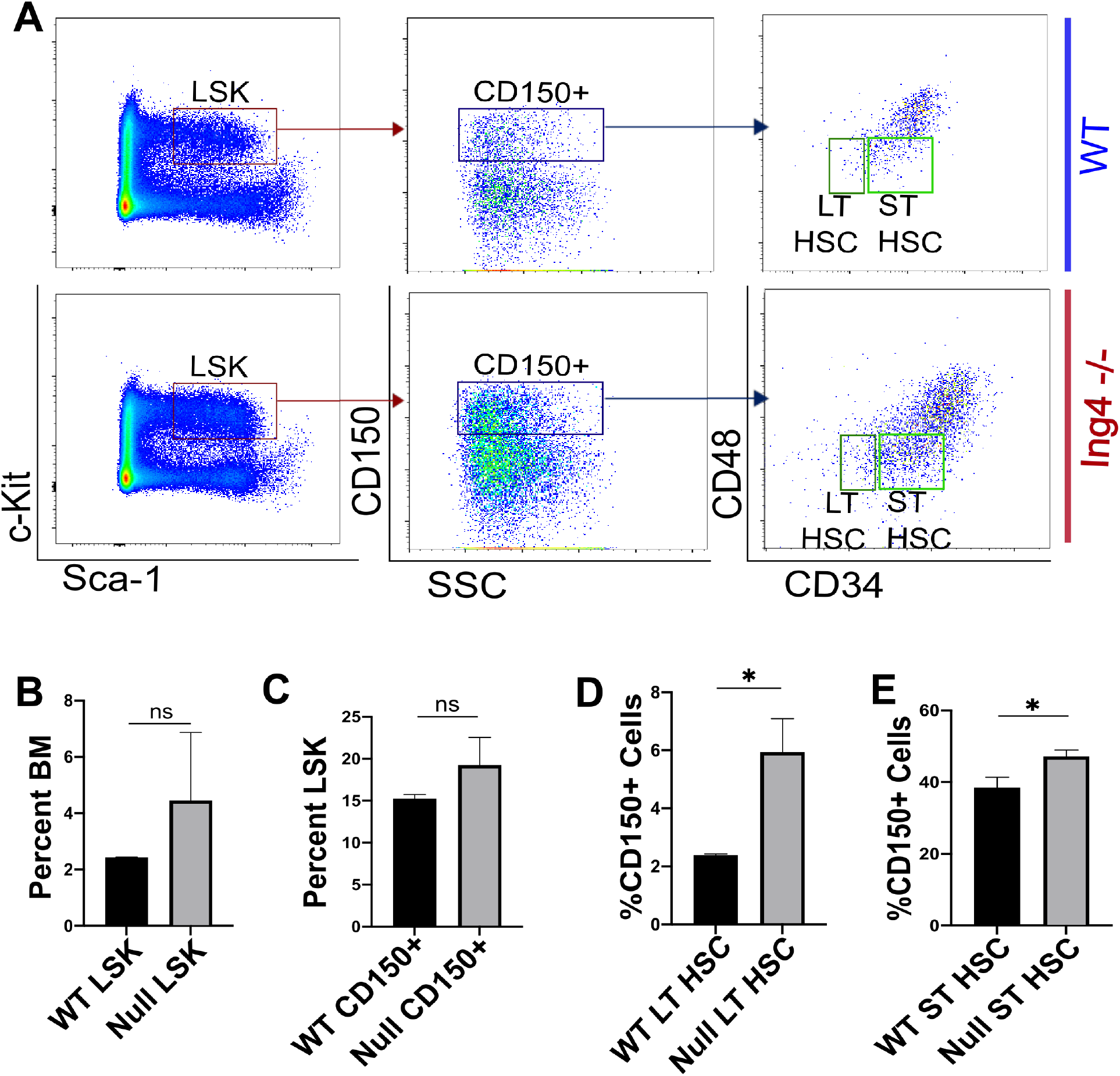
Deletion of Ing4 Leads to Skewed Hematopoiesis in the Bone Marrow. (A) Representative flow cytometric analysis of HSPC subpopulations within the WBM of mice at steady state. (B-E) Frequency of LSK (B), CD150^+^LSK (C), LT-HSC (D), and ST-HSC (E) from the WBM isolated from femora of individual WT and Ing4^−/−^ steady state mice (n = 4-6 per group). ns=p>0.05, *p < 0.05. Data are represented as mean ± SEM.

Analysis via flow cytometry of lineage committed cells in the WBM revealed no significant differences in lineage (Lin^+^) cell populations in the WBM of Ing4^−/−^ mice (Figure 2a). We also analyzed progenitor cell populations in the bone marrow. We found that only the percentage of common lymphoid progenitors (CLPs) was significantly increased in Ing4^−/−^ WBM (Figure 2b,f). These results suggest that Ing4 is largely dispensable for maintenance of the committed progenitor pool.

**Figure 2.**
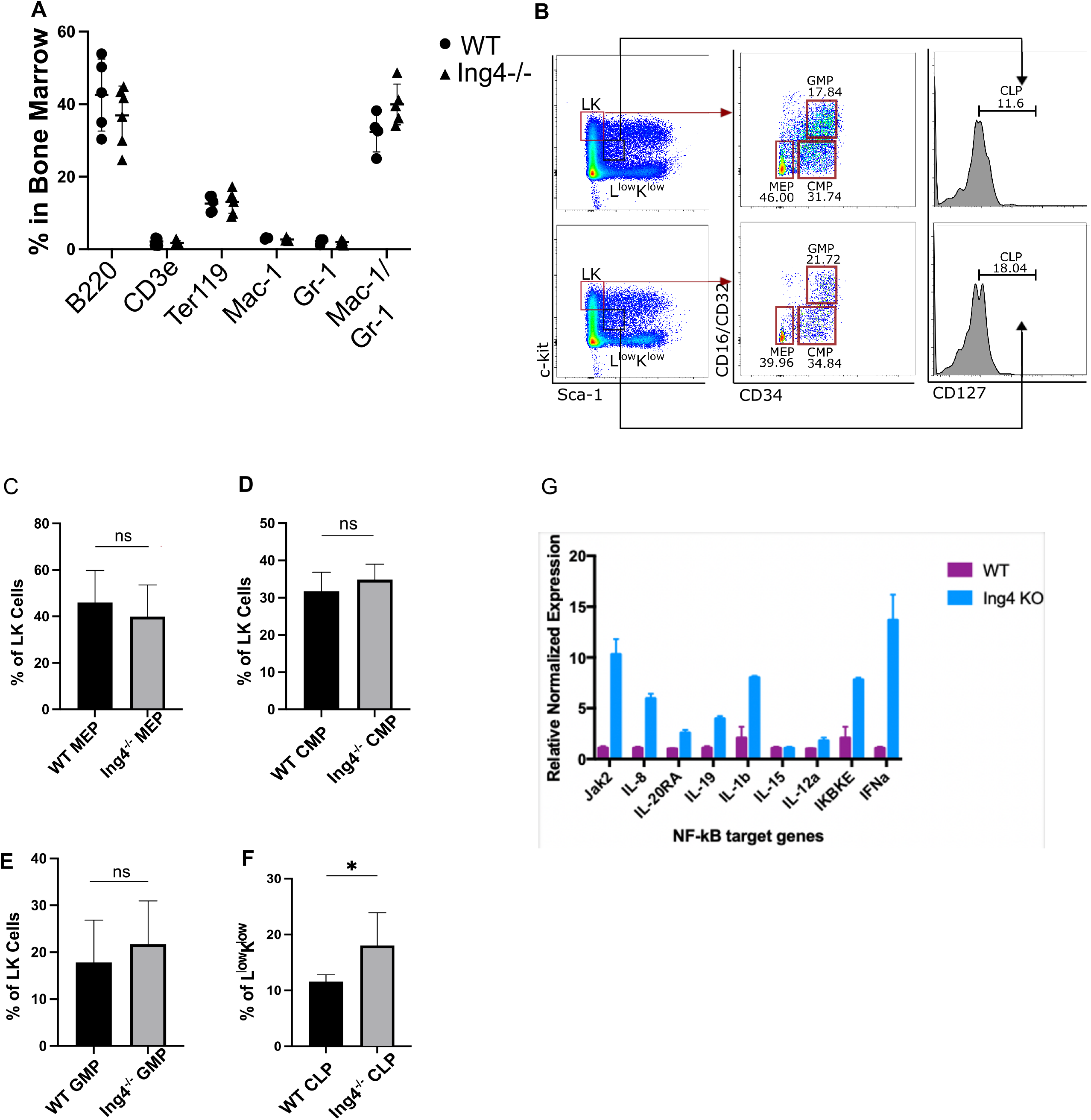
Hematopoietic program is altered in the absence of Ing4. (A) Percentages of lineage committed cell populations in the peripheral blood of WT and Ing4^−/−^ steady-state mice as analyzed via flow cytometry (n=4) *p<0.05, **p<0.01. (B) Percentages of lineage committed cell populations in the WBM of WT and Ing4^−/−^ steady-state mice as analyzed via flow cytometry (n=5). (C) Representative flow cytometry plots of MEP, CMP, GMP, and CLP cell populations in WBM of WT and Ing4^−/−^ steady-state mice. (D) Percentage of MEP cells present in the LK population of WBM cells harvested from WT and Ing4-/- steady-state mice (n=5) ns=p>0.05. (E) Percentage of CMP cells present in the LK population of WBM cells harvested from WT and Ing4^−/−^ steady-state mice (n=5) ns=p>0.05. (F) Percentage of GMP cells present in the LK population of WBM cells harvested from WT and Ing4^−/−^ steady-state mice (n=5) ns=p>0.05. (G) Percentage of MEP cells present in the L^low^K^low^ population of WBM cells harvested from WT and Ing4^−/−^ steady-state mice (n=5) *p<0.05.

Based on previous work demonstrating direct regulation of the p65 subunit of NF-κB by Ing4 and increased cytokine expression in Ing4^−/−^ immune cells, we analyzed cytokine expression in Ing4^−/−^ WBM cells ^23,25^. RT-qPCR analysis of whole bone marrow from Ing4^−/−^ mice confirmed a substantial increase in gene expression patterns of inflammatory cytokines (IL-8, IL-20RA, IL-19, IL-1b, IL-12a, IKBKE, IFNα) in marrow collected from Ing4^−/−^ mice (Figure 2g). These results suggest inflammatory signaling mechanisms are increased in the Ing4^−/−^ bone marrow compartment.

### Ing4 Deficient HSCs Are Quiescent and Have Reduced Intracellular ROS Content

Given the presence of inflammatory cytokines in the bone marrow, we assessed Ing4^−/−^ HSCs for cell cycle status. The balance of quiescence and proliferation are essential for maintaining HSC populations and inflammatory signals typically promote proliferation ^35^. To analyze proliferative status of HSCs, cell cycle analysis was conducted using DAPI and Ki-67. Surprisingly, these experiments revealed while 76.2% of WT LT-HSCs were in G_0_, substantially more LT-HSCs from Ing4^−/−^ mice (92.9%) were detected in G_0_ (Figure 3k,l). ST-HSCs were also more quiescent with 27% of Ing4^−/−^ ST-HSCs in G_0_ compared to 11.1% of wildtype ST-HSCs (Figure 3k,l). To determine if inflammation was causing senescence of HSCs, senescence-associated β galactosidase (SA-β-gal) staining was used. We found a subtle, but statistically significant increase in SA-β-gal activity in Ing4^−/−^ LSK cells (Figure 3d,e). Upon further profiling, we determined that senescence was only increased in the Ing4^−/−^ LT-HSC population (Figure 3f). The low frequency of senescent cells suggest that Ing4 deficiency does not trigger senescence in the majority of quiescent cells.

**Figure 3.**
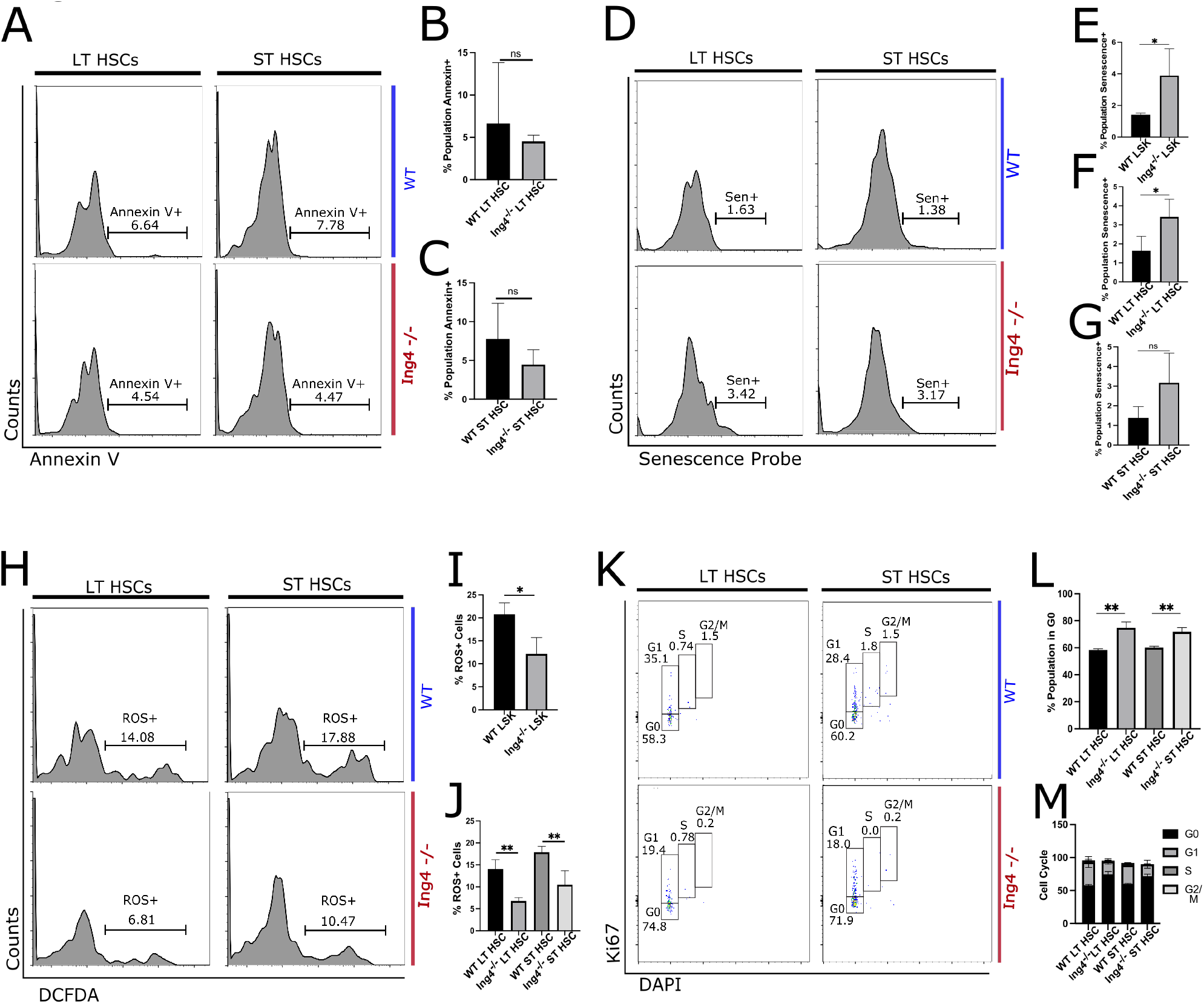
Ing4-deficient HSCs have increased quiescence and senescence. (A) Representative flow cytometry plots of apoptosis analysis of LT (LSK 150^+^34^−^) and ST (LSK150^+^34^+^) HSCs using annexin V. A summary of the percentage of annexin V^+^ cells in LT-HSCs (B) and ST HSC (C) in WT and Ing4^−/−^ marrow at steady-state hematopoiesis (n=5). (D) A representative experiment of percent senescence determination through flow cytometry. A summary of the percentage of senescence cells in LSK (E), LT-HSC (F), and ST-HSC (G) cell populations collected from WT and Ing4^−/−^ WBM at steady-state hematopoiesis (n=4-5) *p<0.5. (H) Representative flow cytometry plots of reactive oxygen species present in LSK cell populations. A summary of ROS^+^ cell percentage in LSK (I), LT-HSC and ST-HSC (J) cell populations in WT and Ing4^−/−^ WBM at steady-stead hematopoiesis (n=5) *p<0.05, **p<0.01. (K) Representative flow cytometry images from cell cycle analysis of LT- and ST-HSCs using Ki-67 and DAPI. (L) Percentage of given cell populations that are in G_0_ of the cell cycle at steadystate hematopoiesis (n=3-4) **p<0.01. (M) Summary of cell cycle analysis of LT-HSC and ST-HSC cell populations in WT and Ing4^−/−^ marrow at steady state hematopoiesis (n=5).

Ing4 has been shown to positively regulate apoptosis through a direct interaction with p53. Absence of Ing4 gives a competitive advantage to tumor cells by limiting p53 activity and, therefore, apoptosis ^36^. We profiled Ing4^−/−^ HSCs for apoptosis to determine if loss of Ing4 enhances evasion of apoptosis in HSCs. When compared with WT HSCs, there was no significant difference in the frequency of cells undergoing apoptosis (annexin V^+^) in the Ing4^−/−^ HSC populations (Figure 3b,c).

Previous studies have shown a direct link between HSC quiescence and low levels of intracellular reactive oxygen species (ROS) content ^37,38^}. To determine ROS levels in Ing4^−/−^ HSCs, we measured freshly isolated LSK cells with 2’,7’-diclorofluoresin diacetate (H2DCFDA) and found that Ing4^−/−^ LSK cells had consistently lower ROS content than WT controls (Figure 3i). Consequently, both Ing4^−/−^ LT-HSC and ST-HSC cell populations had significantly lower ROS content than controls (Figure 3j). These results correlate with Ing4^−/−^ HSC quiescence levels.

### Ing4^−/−^ HSCs Simultaneously Express Genes Associated with Differentiation and Quiescence

To elucidate the molecular consequences of Ing4 loss in HSC regulation, we conducted a genome-wide expression analysis using RNA-seq of purified Ing4^−/−^ and WT LT-HSCs and ST-HSCs. This revealed 2,420 differentially expressed genes (1,139 upregulated and 1,282 downregulated, p<0.05). Gene set enrichment analysis (GSEA) of this data set showed upregulated genes in Ing4^−/−^ HSCs were associated with oxidative phosphorylation and c-Myc target genes and down regulated genes were associated with mitotic spindle formation (Figure 4b-d). Among the most notably upregulated genes, we uncovered genes associated with ribosomal biogenesis, which also correspond with increased c-Myc activity (Figure 4a). Several genes that correlate with HSC quiescence were also upregulated, including cell-cycle regulators p21 and p57, and several genes associated with antioxidant activity (Figure 4a). Together, these signatures suggest that Ing4 deficiency may promote a poised state in HSCs, with upregulation of pathways typically associated with differentiation, yet they retain several quiescence associated genes keeping them arrested in G_0_.

**Figure 4.**
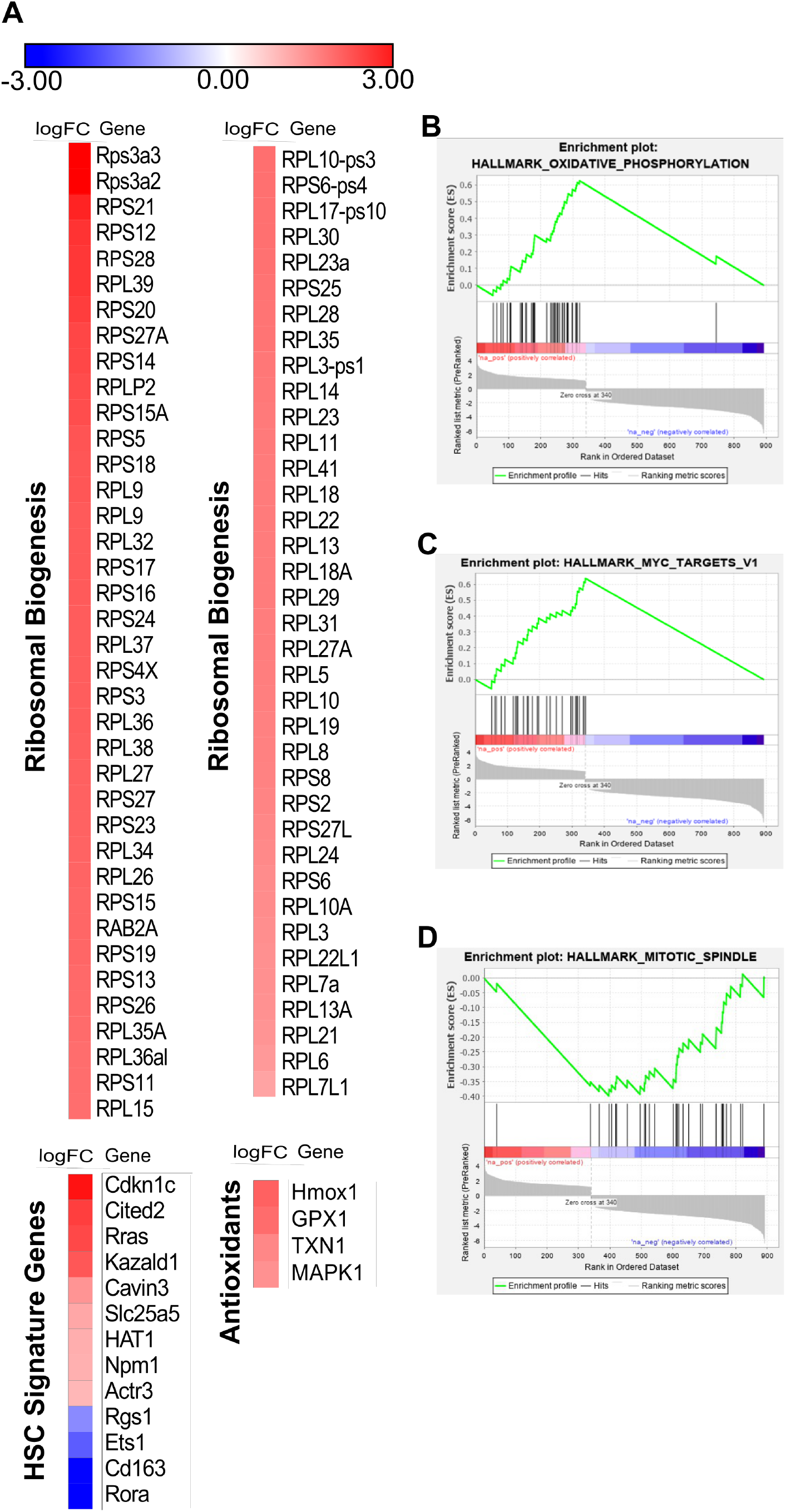
Gene expression analysis of Ing4-/- HSC populations. (A) Heat maps of upregulated genes associated with ribosomal biogenesis, HSC signature, and antioxidants. (B) GSEA plot showing enrichment if upregulated genes in Ing4^−/−^ HSCs for oxidative phosphorylation (B) and myc targets (C), and downregulation of genes associated with mitotic spindle formation (D).

### Ing4 Deficient LT-HSCs Demonstrate Alterations in Stress Hematopoiesis

To analyze the functional role of Ing4 deficiency in HSC reconstitution, we transplanted WBM cells from Ing4^−/−^ and WT mice into lethally (9 Gy) irradiated WT recipients in a competitive setting (1:4 ratio). Peripheral blood analysis at 4, 12, and 16 weeks post-transplantation showed no significant change in donor contribution of Ing4^−/−^ cells in comparison to WT cells (Figure 5c) nor was there a change in lineage distribution observed (data not shown). At 16 weeks post-transplantation, the WBM of engrafted mice was analyzed. This again revealed no significant reduction in chimerism of Ing4^−/−^ HSCs (Figure 5d). These results were striking, given previous models of NF-κB pro-inflammatory cytokine expression, which demonstrated impaired engraftment ^29,30^. Our model demonstrates a perceived ability to overcome this impairment.

**Figure 5.**
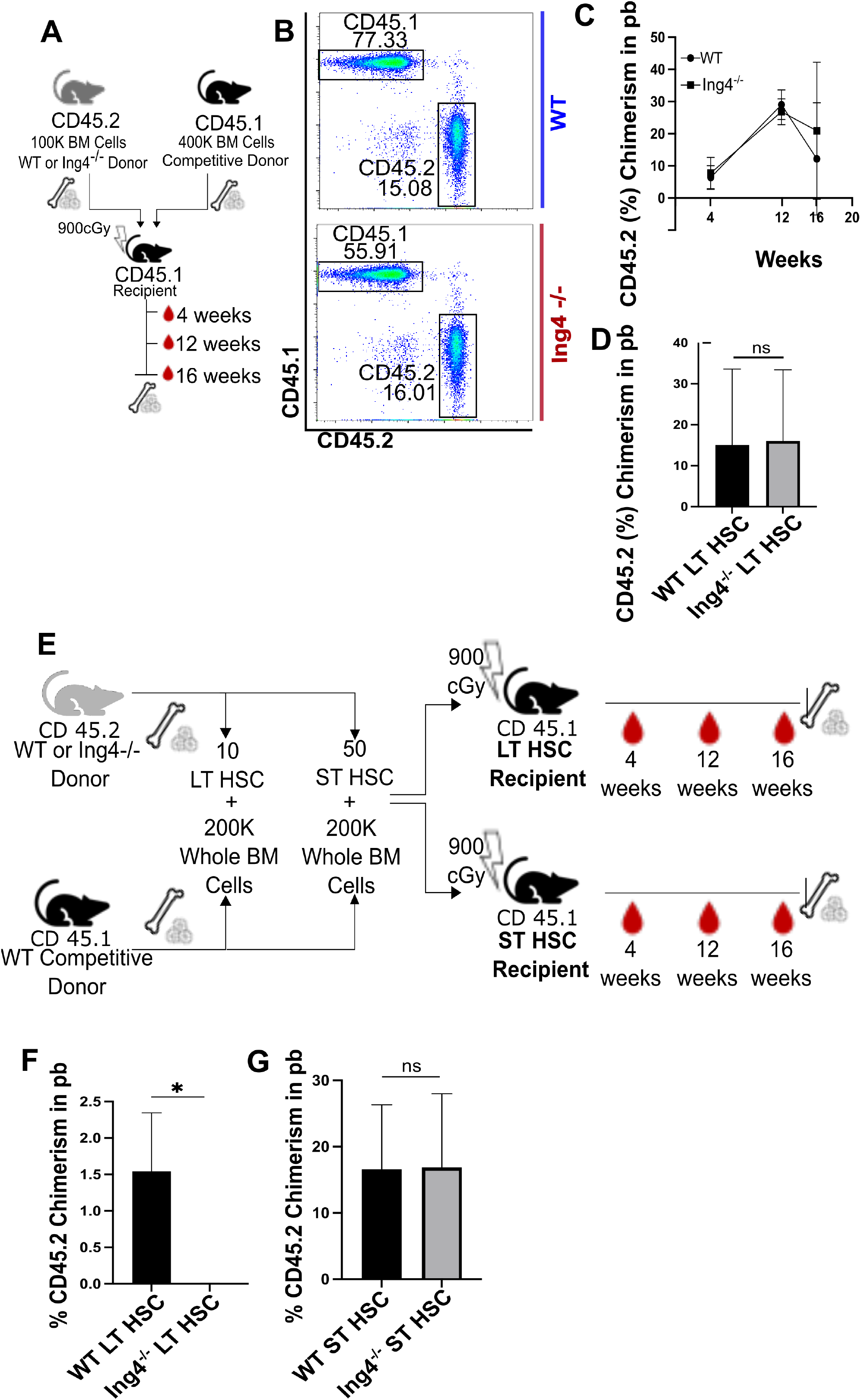
Ing4-/- LT HSCs show alterations in stress hematopoiesis following competitive transplant. (A) Schematic overview of the competitive transplantation assay using WBM from WT and Ing4^−/−^ mice. (B) Representative flow cytometry plots of CD45.1 and CD45.2 chimerism in peripheral blood collected from recipient mice 16 weeks following transplantation with WBM from WT and Ing4^−/−^ mice. (C) Percent CD45.2 chimerism in the peripheral blood of recipient mice collected 4, 12, and 16 weeks following competitive WBM transplantation (n=5). (D) Percent CD45.3 chimerism in the peripheral blood of recipient mice collected 16 weeks post competitive WBM transplantation (n=5) ns=p>0.05. (E) Schematic overview of competitive transplantation assay using sorted LT- and ST-HSCs from WT and Ing4^−/−^ mice. (F) Percent CD45.2 chimerism in the peripheral blood of recipient mice collected at 16 weeks post competitive transplant with LT-HSCs from WT and Ing4^−/−^ donor mice (n=10) *p<0.05. (G) Percent CD45.2 chimerism in the peripheral blood of recipient mice collected at 16 weeks post competitive transplant with ST HSCs from WT and Ing4^−/−^ donor mice (n=10) ns=p>0.05.

To more directly establish the reconstitution and maintenance capacity of Ing4 deficient HSC subsets, we transplanted sorted LT-HSCs and ST-HSCs from CD45.2 Ing4^−/−^ and WT mice in a competitive setting, 10 LT-HSCs or 50 ST-HSCs combined with 200,000 CD45.1 unfractionated WBM cells. PB analysis at 4, 12, and 16 weeks posttransplantation revealed a significant reduction in donor contribution of the Ing4^−/−^ LT-HSCs in comparison to WT LT-HSC (Figure 5f.), however there were no significant differences in the chimerism observed in ST-HSC populations (Figure 5g). These results suggest Ing4^−/−^ LT-HSCs are incapable of repopulation, but ST-HSCs have robust repopulating activity. The potential effects of inflammation may impair the function of Ing4^−/−^ LT-HSCs, but not ST-HSCs ^39^, which are likely responsible for the engraftment observed in our WBM transplantation.

To further dissect hematopoietic function in the absence of Ing4, we studied the response of Ing4^−/−^ HSCs to hematopoietic stress induced by myeloablative agent 5-fluorouracil (5-FU) as a mechanism to trigger HSC activation. Following single dose 5-FU injection, there was a significant decrease in Ing4^−/−^ LSK population, that unlike the WT LSKs, failed to recover (Figure 6b). Both Ing4^−/−^ LT-and ST-HSC populations failed to recover up to 14 days following 5-FU treatment indicating a reduced sensitivity of 5-FU ablation (Figure 6d,e). These results were unexpected as we hypothesized that ST-HSCs would recover from myeloablation, however, 5-FU has previously been shown to alter ribosomal biogenesis in addition to targeting DNA ^40–42^. Thus, 5-FU may be impairing one of the major mis-regulated pathways in Ing4^−/−^ cells in addition to inducing DNA damage. To determine whether the more severe WBM failure induced by 5-FU treatment was due to these off-target effects, we used sublethal irradiation (3.5Gy) as an alternative insult to target dividing cells. 14 days post-irradiation, Ing4^−/−^ mice showed a significant reduction in LT-HSC population, however Ing4^−/−^ ST-HSCs exhibited recovery from sublethal irradiation similar to that of WT ST-HSCs (Figure 6i,j). These results show that DNA damage activates Ing4^−/−^ ST-HSCs, but not LT-HSCs, however this activation can be impaired by the off-target effects of 5-FU.

**Figure 6.**
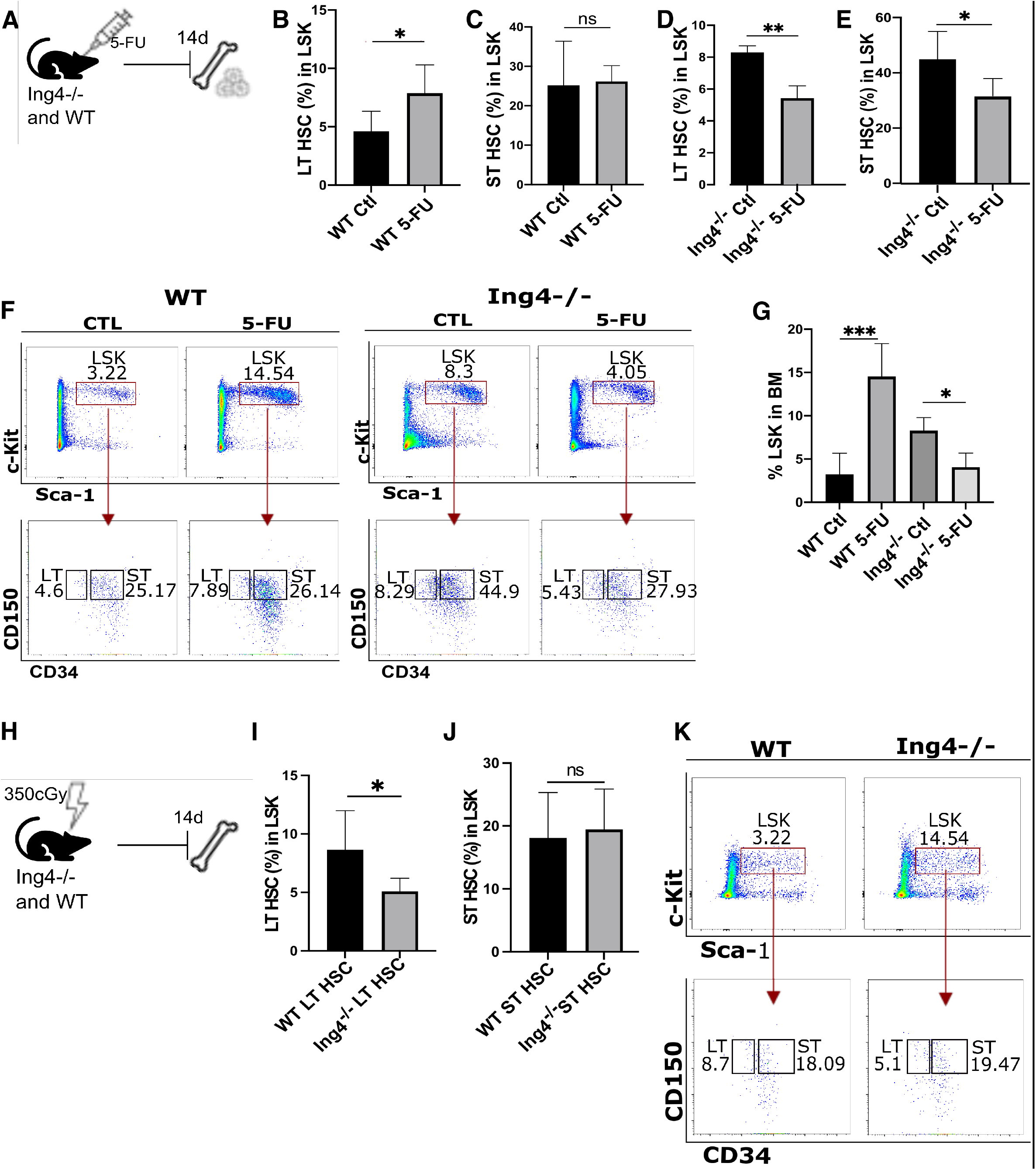
Ing4-Deficient HSCs show alterations in stress hematopoiesis following chemotoxic insult and irradiation. (A) Schematic representation of 5-FU treatment of steady-state mice. (B) Percentage of LT-HSCs present in LSK cell population in WBM of WT steady-state mice following one treatment of 5-FU (n=5) or PBS as a control (n=6) *p<0.05. (C) Percentage of ST-HSCs present in LSK cell population in WBM of WT steady-state mice following one treatment of 5-FU (n=5) or PBS as a control (n=6) ns=p>0.05. (D) Percentage of LT-HSCs present in LSK cell population in WBM of Ing4^−/−^ steady-state mice following one treatment of 5-FU (n=5) or PBS as a control (n=4) **p<0.01. (E) Percentage of LT-HSCs present in LSK cell population in WBM of WT steady-state mice following one treatment of 5-FU (n=5) or PBS as a control (n=4) *p<0.05. (F) Representative flow cytometry plots of LSK cells, LT-HSCs and ST-HSCs in WBM of Ing4^−/−^ and WT steady-state mice following treatment with 5-FU. (G) Percentage of LS cells in WBM 14 days after 5-FU administration (n=4-6) *p<0.05, ***p<0.001. (H) Schematic representation of the experimental strategy for evaluating the effects of sublethal irradiation of steady-state mice. (I) Percentage of LT-HSCs present in LSK cell population in WBM of WT (n=6) and Ing4^−/−^ (n=8) steady-state mice 14 days after sublethal irradiation *p<0.05. (J) Percentage of ST-HSCs present in LSK cell population in WBM of WT and Ing4^−/−^ steady-state mice 14 days after sublethal irradiation (n=10) ns=p>0.05.

### c-Myc Inhibitor Enhances HSC Recovery

We sought to target the mis-regulated pathways in Ing4^−/−^ LT-HSCs to rescue their loss of function. We chose to target the c-Myc pathway for several reasons; c-Myc target genes are over represented in our RNA-seq data and c-Myc lies upstream of several of the other mis-regulated pathways observed in Ing4^−/−^ HSCs. Finally, Ing4 has previously be reported to negatively regulate c-Myc activity ^26^. c-Myc, however, has a controversial history in hematopoiesis. Previous studies have shown that c-Myc inactivation results in an accumulation of HSCs and diminished HSC function ^43,44^, but also can enhance HSC expansion in culture ^45^. Enforced expression of c-Myc results in HSC loss of function, at least in part through N-cadherin and integrin down regulation ^44,46^. Yet, increased c-Myc expression has been shown to promote HSC self-renewal ^47^. Specific levels of c-Myc protein appear to be critically important for HSCs, perhaps resulting in some of the disparate data in the literature ^46,48^.

In Ing4^−/−^ HSCs, c-Myc target genes are highly overexpressed. To ascertain whether Ing4^−/−^ HSC repopulation capacity could be rescued through c-Myc inhibition, we conducted an *in vivo* treatment of Ing4^−/−^ and WT control mice with c-Myc dimerization inhibitor, 10058-F4, following hematopoietic insult by sublethal irradiation. We found that the treatment stimulated both LT-HSCs and ST-HSCs into cycling, but resulted in fewer total LT- and ST-HSCs (Figure 7b,c.). Cell cycle analysis of c-Myc inhibitor treated HSCs revealed that the percentages of Ing4^−/−^ LT- and ST-HSCs in the G_0_ phase of the cell cycle decreased when compared to untreated Ing4^−/−^ HSCs (Figure 7f,h). Consequently, the population of Ing4^−/−^ LT-HSCs in the G_1_ phase of the cell cycle increased indicating a release from G_0_ arrest upon drug treatment (Figure 7g). The population of Ing4^−/−^ ST-HSCs in the S phase of the cell cycle presented a significant increase also indicating release from G_0_ arrest following drug treatment (Figure 7i).

**Figure 7.**
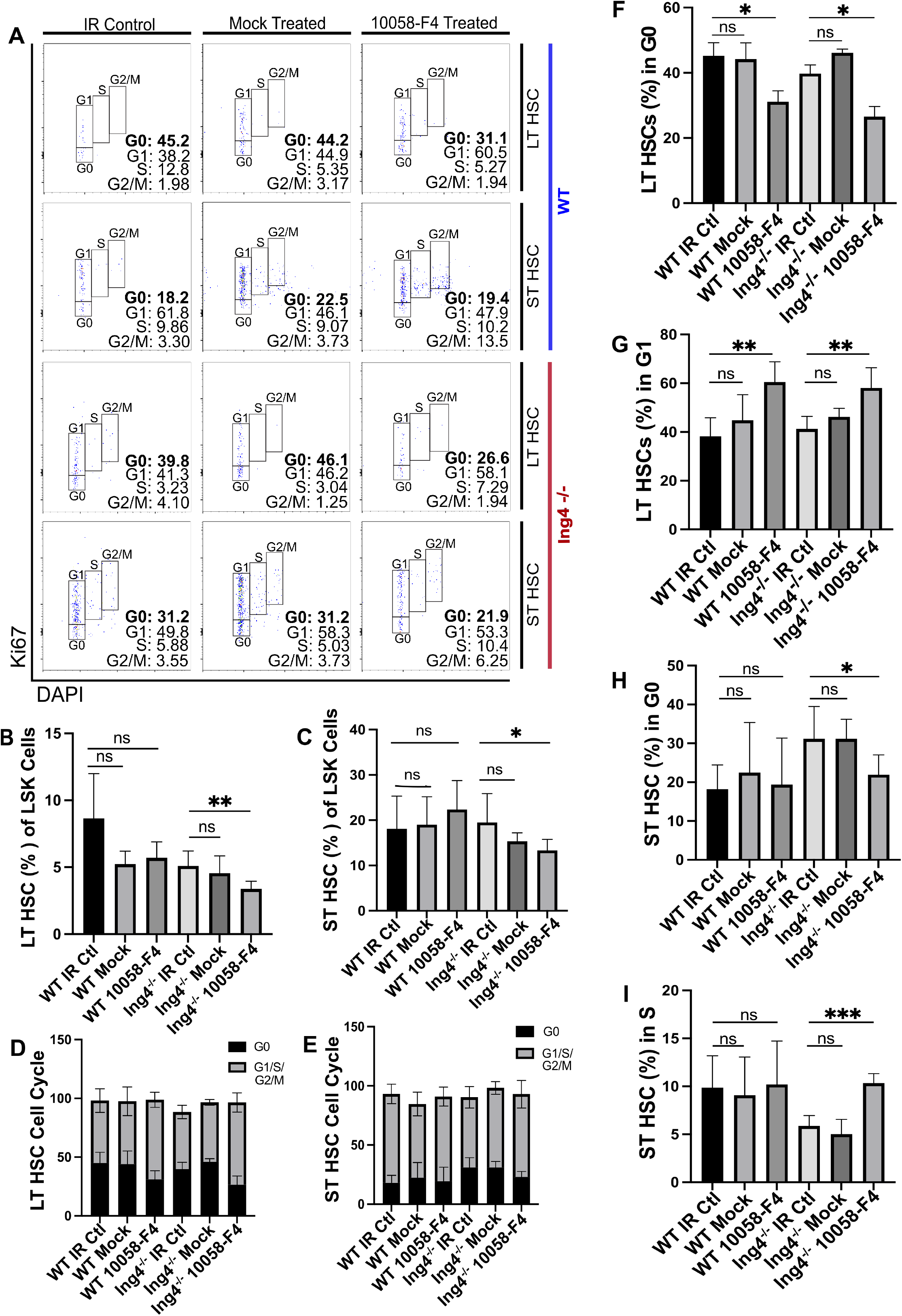
c-Myc inhibition enhances HSC recovery. (A) Representative flow cytometry plots of LT-HSCs and ST-HSCs in WBM of Ing4^−/−^ and WT mice following sublethal irradiation and 14 days of treatment with 10058-F4 or mock treatment with vehicle. (B) Percentages of LT-HSCs present in the LSK populations of cells collected from WBM of WT and Ing4^−/−^ mice following sublethal irradiation, treatment with mock vehicle, and/or 10058-F4 (n=5) ns=p>0.05, **p<0.01. (C) Percentages of ST-HSCs present in the LSK populations of cells collected from WBM of WT and Ing4^−/−^ mice following sublethal irradiation, treatment with mock vehicle, and/or 10058-F4 (n=5) ns=p>0.05, *p<0.05. (D) Summary of cell cycle analysis conducted using Ki-67 and DAPI of LT-HSCs collected from WBM of WT and Ing4^−/−^ mice following 10058-F4 or vehicle treatment. (E) Summary of cell cycle analysis conducted using Ki-67 and DAPI of ST-HSCs collected from WBM of WT and Ing4^−/−^ mice following 10058-F4 or vehicle treatment. (F) Percentages of LT-HSCs in the G_0_ phase of cell cycle is specific treatment groups (n=5) *p<0.05. (G) Percentages of LT-HSCs in the G_1_ phase of cell cycle is specific treatment groups (n=5) **p<0.01. (H) Percentages of ST-HSCs in the G_0_ phase of cell cycle is specific treatment groups (n=5) ns=p>0.05, *p<0.05. (I) Percentages of ST-HSCs in the S phase of cell cycle is specific treatment groups (n=5) ns=p>0.05, ***p<0.001.

## Discussion

The maintenance of HSC homeostasis in the bone marrow, allowing for quiescence, selfrenewal, and differentiation, remains a poorly characterized process. In this study, we elucidate the role of Ing4 in HSC quiescence and self-renewal. Previous work in cancer models has identified Ing4 as a tumor suppressor protein that negatively regulates NF-κB, c-Myc, and HIF-1α while positively regulating p53 and histone acetylation through the Hbo1 complex. Ing4 deficiency in tumor cells provides an environment of increased inflammation, increased c-Myc activity, and decreased p53 activity. Prior to our work, the role of Ing4 in hematopoiesis had yet to be understood. Here, we show that Ing4 plays a critical role as a vital regulator of HSC quiescence and self-renewal.

Ing4 deficiency results in skewed hematopoiesis, with a subtle increase in LT-HSCs, and a large increase in ST-HSCs. Bone marrow cells in Ing4^−/−^ mice express high levels of inflammatory cytokines. Precise regulation of NF-κB signaling in hematopoiesis is critical and skewing of NF-κB activity in either direction has significant consequences ^49–58,59,60,29^. Despite enhanced NF-κB activity, whole bone marrow from Ing4^−/−^ mice shows no difference in repopulation in bone marrow transplantation. This was unexpected given other inflammation models, where whole bone marrow cannot repopulate in transplantation, shown both with genetic and disease models ^29,30,51,55–57,61,62^. Our results suggest that Ing4 deficiency gives rise to regenerative capabilities in HSCs despite the presence of inflammatory signals, conferring resistance to inflammatory stress.

When subpopulations of HSCs are transplanted, the Ing4^−/−^ LT-HSCs show impaired repopulation ability, while ST-HSCs show activity indistinguishable from wild-type ST-HSCs. To understand why ST-HSCs are capable of repopulation despite increased inflammatory signals, but LT-HSCs are not, we analyzed the cell cycle profiles of the HSCs. Both LT-HSCs and ST-HSCs are more quiescent than their wild-type counterparts. Previous models of inflammation have shown that chronic exposure to inflammatory signals results in HSC quiescence as a protective measure ^39^. Furthermore, in dnmt3-deficient HSCs, LT-HSCs proliferate to maintain the stem cell pool rather than for repopulation ^63^. Thus, our data suggest that Ing4^−/−^ HSCs are unable to overcome inflammatory stress and remain quiescent but may proliferate to maintain the stem cell pool. ST-HSCs are able to overcome this barrier.

To identify how ST-HSCs from Ing4^−/−^ mice may be protected from inflammatory stress, we analyzed RNA expression profiles. Markers of oxidative phosphorylation, ribosomal biogenesis, and other c-Myc target genes are significantly upregulated in Ing4^−/−^ HSCs, more closely matching profiles of cycling or differentiating HSCs ^64,65^. Ing4 has previously been identified as a regulator of c-Myc and ribosomal biogenesis ^26,66^. Our results correlate with these associations. Despite these upregulated pathways, Ing4^−/−^ HSCs retain high levels of several stem cell quiescence markers, including Egr1, Bmi1, p57, Gata2, Npm1, and Hat1. These genes have previously been show to play a role in HSC quiescence, providing a mechanism for sustained G_0_ arrest ^67–75^. Together, these signatures suggest that Ing4-deficiency triggers a differentiation program, but cells remain poised for expansion through retention of quiescence. For Ing4^−/−^ ST-HSCs, this results in a competitive advantage in an inflammatory environment that is normally hostile to emergency HSC function.

To further challenge the HSCs, we performed low-dose irradiation treatment. These assays promote HSC response by inducing damage in differentiated immune cells, which require repopulation from HSCs. As in our transplantation assay, Ing4^−/−^ LT-HSCs are unable to respond to this challenge, but ST-HSCs repopulate at wild-type levels. To trigger proliferation and overcome the mis-regulated pathways in Ing4^−/−^ HSCs, we used a c-Myc inhibitor to target several of the pathways overexpressed in our RNA-seq analysis, including ribosomal biogenesis, oxidative phosphorylation, and c-Myc target genes. This inhibitor triggered proliferation in both LT-HSCs and ST-HSCs, but resulted in a reduction of cells in these populations. These results suggesting that Ing4 deficiency can be partially overcome through targeting of c-Myc associated pathways, but is accompanied by a loss of function, likely through apoptosis ^39^.

Our results demonstrate for the first time the importance of Ing4 regulation of several pathways required for hematopoiesis and shed new light on mechanisms for resistance to inflammatory stresses. Ing4 lies upstream of several pathways that regulate stem cell quiescence and differentiation. Our results suggest that manipulation of these pathways improve the function of ST-HSCs, placing them in a quiescent, but poised state that can overcome inflammatory stress. Increased ribosomal biogenesis and other c-Myc associated genes may be involved, as c-Myc inhibition can impair this resistance to inflammatory stress, but inhibition of c-Myc has many outcomes. Further studies are necessary to elucidate which specific pathways downstream of Ing4 are important for conferring the inflammatory resistance in Ing4 deficiency. Manipulation of these pathways may provide a mechanism to improve HSC function during hematopoietic stress.

## Acknowledgements

This work was funded by NIH K01DK104974-01 and NIH 5P20GM109091-05. We would like to thank the Flow Cytometry Core Facilities at the University of South Carolina (UofSC) School of Pharmacy. Specifically, we thank Chang-Uk Lim for his assistance with flow cytometry sorting, analysis, and experimental design. We also thank Jason Kubinak and Kia Zellars for their assistance in using the Instrumentation Resource Facility flow cytometer at the UofSC School of Medicine. For animal support, we thank the University of South Carolina Department of Laboratory Animal Research core members Shaun Thacker and Marlee Poole.

## Author Contributions

Conceptualization, KLK, ZT, Methodology, KLK, ZT, GA, MR, SG, VB, Data Analysis, KLK, ZT, VB, AMT, Writing, KLK, ZT, Funding Acquisition, KLK, Supervision, KLK.

## Materials and methods

### Animals

Ing4^−/−^ mice (CD45.2) were provided by Stephen N. Jones (University of Massachusetts Medical School) and mice were backcrossed to C57BL/6 on site. Wildtype (WT) C57BL/6 (CD45.2) and C57Bl/6.SJL (CD45.1) mice 8-12 weeks of age served as controls. C57BL/6 (CD45.2) and B6. SJL-Ptprc^a^ Pepc^b^/BoyJ (CD45.1) were purchased from The Jackson Laboratory. All mice were maintained at an AALAC-accredited animal facility at the University of South Carolina according to Institutional Animal Care and Use Committee animal guidelines, and all animal experiments were performed with consent from the local ethical committee (protocol #101536). Genotypes were confirmed by PCR of genomic DNA.

### Quantitative real-time PCR and cytokine detection

For quantitative RT-PCR (qRT-PCR) in zebrafish, total RNA from whole embryos was extracted with TRIzol (Invitrogen). cDNA was prepared with SuperScript III First Strand Synthesis System for RT-PCR (Invitrogen) using oligo-dT primers. qRT-PCR was performed using the ZYBR Green PCR Master Mix (Applied Biosystems). Gene expression changes were quantified as linear ration to β-actin.

For qRT-PCR in mice, total RNA from bone marrow cells was extracted with TRIzol (Invitrogen). (insert probes) probes were used with Superscript III for RT-PCR (Invitrogen) using oligo-dT primers. Triplicate samples were normalized to β-actin and Gapdh controls.

IFN-γ, TNF-α, IL-6, IL-10, MCP-1, and IL-12p70 levels in mouse bone marrow supernatant were determined using BD Cytometric Bead Array Mouse Inflammation Kit. Bone marrow supernatant was isolated from mouse tibias and femurs by suspending the bones in p200 tips trimmed to fit into 1.5 ml Eppendorf tubes filled with 1 ml PBS. Tubes were centrifuged for 8 min at 500g. Marrow was centrifuged again, and the clear supernatant was isolated. The protein content of the supernatant was quantified via a NanoDrop spectrophotometer (NanoDrop Technologies).

### Cell sorting and transplantation assays

For HSC isolation (Lin^−^Sca^−^c-Kit^+^[LSK]CD48^−^CD150^+^ cell fraction), whole bone marrow (WBM) cells isolated by crushing femur and tibia from each mouse were incubated with 1x RBC Lysis Solution (Miltenyi Biotec) to eliminate red blood cells. Cells were lineage depleted using a Lineage Cell Deptletion Kit (Miltenyi Biotec) and passed through a nylon filter (40um; Fischer) to obtain single-cell suspensions followed by cell sorting.

For WBM competitive cell transplantation, 1 × 10^5^ total WBM cells were isolated from Ing4^−/−^ mice and control donor mice (all on CD45.2 background) and transplanted with 4 × 10^5^ CD45.1+ competitor cells via retro orbital (r.o.) injection into 8-to 12-week-old recipient mice (B6. SJL-CD45.1) that had been lethally irradiated (9 Gy split in 2 doses, at least 3 hours apart).

For competitive transplantation assays, 10 LT-HSCs or 50 ST-HSCs were sorted from Ing4^−/−^ or WT donors (CD45.2) were combined with 200,000 CD45.1 total, unfractionated WBM cells and transplanted into sublethally irradiated recipient mice (CD45.1). Flow cytometry analysis was performed using a BD LSRII and sorting was performed using a BD FACSAria II cytometer. Data was analyzed using FlowJo version 10.6.2 software. Transplantation strategies are outline in Figure 5, and antibody combination is provided in supplemental Table 1.

### Peripheral blood analysis

Peripheral blood collected via r.o. vein was collected using heparinized micro-hematocrit capillary tubes (Kimble) and kept in heparin (0.05 IU/mL; Alfa Aesar). Before antibody staining, erythrocytes were lysed with 1X red blood cell lysis solution (Miltenyi Biotec) for 10 minutes at room temperature. Cells were stained with antibodies against CD45.2 and CD45.1 to assess engraftments levels, as well as antibodies against lineage positive markers CD3e (T cells), CD45R (B cells), Ly-6G/C (granulocytes), Ter119 (erythrocytes), and Mac-1 (macrophages) to determine donor contribution as well as lineage skewing.

### Cell differentiation analysis

WBM cells were incubated for 30 minutes at 4°C with antibodies for HSC isolation (CD34, CD48, Sca, c-Kit; Invitrogen), lineage markers (CD3e, CD45R, Ter119, Ly-6G/C, and CD11b; Invitrogen) as well as anti-mouse CD16/32 and CD127 (Biolegend). Cells were washed twice with phosphate-buffered saline (PBS)-bovine serum albumin (BSA).

### Cell cycle and cell proliferation analysis

WBM cells were incubated for 30 minutes at 4°C with antibodies for HSC isolation. Cells were washed with PBS-BSA and fixed and permeabolized according to manufacturer’s instructions. Cells were incubated in the dark for 30 minutes at room temperature in Click-iT reaction cocktail. Next, cells were incubated for 15 minutes at room temperature in 50 ug/mL propidium iodide (Invitrogen), washed with PBS-BSA, and analyzed using a BD LSRII.

### 5-Fluorouracil treatment and Sublethal Irradiation

8-to 12-week-old Ing4^−/−^ and WT mice were r.o. injected once with 5-fluorouracil (5-FU; 150 mg/kg each). 14 days following 5-FU injection, WBM was harvested, lineage depleted, and analyzed via flow cytometry. To give a sublethal dose of radiation, mice were irradiated with 3.5 Gy total body irradiation

### c-Myc Inhibitor Treatment

8-to 12-week-old Ing4^−/−^ and WT mice were irradiated with a sublethal dose (3.5 Gy) of total body irradiation. One day after irradiation, mice began intraperiteal (i.p.) injections of 25mg/kg/day of dimerization inhibitor, 10058-F4 (Cayman) dissolved in DMSO and diluted with corn oil or a mock treatment with DMSO as a control. Following 14 days of treatment, bone marrow was harvested for flow cytometry analysis.

### RNA-seq analysis

Approximately x HSCs (LSK^+^CD150^+^) were sorted into Buffer RLT Plus from pools of WT or Ing4^−/−^ mice. RNA was isolated with the RNeasy Micro column (QIAGEN). Illumina HiSeq was used for sequencing…

### Flow Cytometry

For lineage analysis, cells were stained with the combinations of fluorochrome-conjugated monoclonal antibodies for cell-surface markers for 30 min in darkness at 4°C. For stem and progenitor cell analysis and sorting purposes, WBM cells were lineage depleted using anti-lineage microbeads with the MACS separation column (Miltenyi Biotec) according to manufacturing protocol, followed by staining with the combinations of fluorochrome-conjugated monoclonal antibodies.

### Measurement of ROS

Red-cell lysed WBM cells were incubated with 7.5uM H2DCFDA (company) resuspended in PBS or PBS alone for unstained controls for 30 minutes at 37 degrees C. The cells were washed with PBS and resuspended in PBS for immediate analysis on a FACSLSRII, and MFI was compared with unstained control cells.

### Measure of Senescence

Lineage depleted bone marrow cells were stained with antibodies to label surface markers, then fixed with the manufacturer provided fixation solution. Fixed cells were incubated with CellEvent Senescence Probe (Thermo) for 2 hours at 37°C with CO_2_, washed, and analyzed by flow cytometry.

### Statistics

Mean values ± SEM are shown. Student’s t test or two-way ANOVA were used for comparisons (GraphPad Prism v.9.0.0). *P<0.05, **P<0.01, ***P<0.001.

See Supplemental Experimental Procedures for more details.

